# Temperate Phage Shape Honey Bee Gut Microbiome Structure and Response to Antibiotic Treatment

**DOI:** 10.64898/2026.06.12.731954

**Authors:** Dino Lorenzo Sbardellati, Arin Mehra, Vincent A. Ricigliano, Julia Fine, Rachel Lee Vannette

## Abstract

Bacteriophage (phage) are hypothesized to play a significant role in modulating gut microbiomes. Yet, in vivo research examining the role of phage in gut ecosystems remains sparse, largely due to a lack of tractable model systems. Here, we use the honey bee (Apis mellifera) gut as a model to test the hypotheses that phage-bacteria interactions in the gut are temporally variable and that stress, in the form of antibiotics, can enhance phage killing of their bacterial hosts. First, we isolated and characterized a novel temperate phage which infects a bee-specific strain of Bifidobacterium. We then mono-colonized adult honey bees with this Bifidobacterium strain and tested how phage treatment impacts bacterial abundance over time. Next, using a series of in vitro and in vivo experiments, we examined how phage-bacteria interactions change in response to treatment with tetracycline, an antibiotic commonly used in commercial beekeeping. Finally, to probe the biological mechanisms underlying different phage-antibiotic synergies, we assayed how different classes of antibiotics impacted bacterial growth with and without phage infection. Our results indicate that a single temperate phage can both promote the ability of its Bifidobacterium host to colonize the gut, while also increasing host sensitivity to antibiotic treatment. Together, these findings demonstrate that environmental stressors can shift phage-bacteria interactions from mutualism to antagonism, highlighting the significance of phage in shaping how gut microbiomes respond to antibiotic or xenobiotic perturbation.

## Introduction

Bacteriophage (phage), the viruses which infect bacteria, are an essential component of gut microbiomes^1–3^. Phage can directly shape microbiome composition by lysing dominant bacteria and facilitating the establishment of otherwise competitively excluded microbes^4,5^. Beyond predation, phage can also influence bacterial communities by mediating the horizontal transfer of genes which alter bacterial metabolism, physiology, and competitive dynamics^6,7^. Despite their potential importance, phage are often overlooked in basic gut microbiology studies. In particular, there is a lack of experimental work examining how different antibiotics influence phage-bacteria interactions in the gut. Antibiotics represent a powerful, tractable, and relevant tool for probing microbiomes, as they can impose mechanistically distinct forms of stress on microbial communities and are nearly ubiquitous in modern healthcare and agriculture^8^. Using antibiotics to build a more complete understanding of how stress and phage interact to shape gut microbiomes will improve our understanding of the basic forces shaping gut microbiology and will enhance our ability to predict how gut microbiomes respond to different forms of stress, including medical interventions.

Antibiotic application is well understood to disrupt gut microbiomes^8–10^ and numerous studies have shown that phage and antibiotics can act synergistically to inhibit or kill bacteria^11–13^. However, until recently, these two phenomena have largely been studied in isolation, with only a few studies examining how phage-antibiotic synergy contributes to gut microbiome disruption by antibiotics. Using metagenomics, Pfeifer et al.^14^ showed that antibiotic treatment can substantially alter phage–bacteria interactions in the human gut. Similarly, Franklin et al.^15^ leveraged a controlled mouse model to demonstrate that the presence or absence of phage communities can modulate bacterial sensitivity to antibiotics. While these studies have advanced our understanding of how phage contribute to microbiome stress responses, there remain gaps in our knowledge. Namely, these studies focus primarily on virulent phage and rely on undefined phage consortia, rather than isolated phage-bacteria pairs. This limits our ability to resolve how both temperate phage and individual phage-bacteria pairs respond to antibiotic stress.

Phage can be broadly categorized as virulent or temperate based on their life cycles. Virulent phage are obligately lytic and rapidly replicate and kill their host following infection, whereas temperate phage generally have the ability to integrate into host genomes following infection, where they can remain dormant for generations^1^. Temperate phage dominate the gut microbiomes of humans and other animals^16^, with an estimated >70% of all gut bacteria harboring one or more of these viruses^17^. Importantly, virulent and temperate phages likely respond differently to antibiotic stress. Virulent phage–antibiotic synergy has been shown to be driven by increased viral absorption^18^ and virion production^19^, whereas temperate phage-antibiotic synergy can occur through biasing lysis-lysogeny decision making^20^ and by inducing dormant phage^21^. Consequently, antibiotics and other forms of stress, may differentially alter phage–bacteria dynamics in the gut depending on phage lifestyle. Given the dominance of temperate phage in the gut, and the widespread use of antibiotics in modern medicine and agriculture^22,23^, understanding how temperate phage interact with commonly used antibiotics will be critical for predicting, and potentially guiding, how microbiomes respond to stress.

A central barrier to studying phage-bacteria interactions in the gut is the sheer complexity typically associated with gut microbiomes. The human gut microbiome is highly diverse and individualized^24^, containing hundreds of different bacterial taxa, each of which is host to its own unique collection of phage^25,26^. The bee gut microbiome represents an ideal model for overcoming this obstacle. Social bees house a simple (5-9 taxa), highly conserved, and well-characterized gut bacterial community^27,28^. Yet, like humans and other animals, bees rely on this simplified gut bacterial community for essential functions, including pathogen defense and nutrient acquisition^29–31^. The bee gut microbiome is also highly tractable, with microbiota depleted and gnotobiotic animals readily reared in the lab^27,32^. Recently, metagenomic surveys have established that honey bees house phage and other mobile genetic elements which target members of their highly conserved gut bacterial community^33–38^. However, the functional role of these phages in shaping gut ecology is less well understood. In particular, how phage-bacteria interactions in the bee gut respond to antibiotic stress remains largely untested. This is especially relevant as commercially managed honey bees are routinely treated with antibiotics^39^. Developing a better understanding of phage-bacteria interactions in the bee gut will help us better understand the importance of phage in other gut communities, while also yielding insights into honey bee management decisions and pollinator health^40^.

Here, we use honey bees as a model system to test how infection with a temperate phage changes the *in vitro* and *in vivo* ecology of *Bifidobacterium*, a prominent beneficial bacterium and important bee gut symbiont. We hypothesize that phage treatment will reduce *Bifidobacterium* growth and abundance and that biological stress, in the form of antibiotic treatment, will enhance this negative impact. To test this hypothesis, we first isolate a complex-carbohydrate degrading *Bifidobacterium* from the bee gut, as well as a novel phage which infects this bacterium. We then colonize microbiota-depleted bees with this phage-bacteria pair and monitor bacterial establishment and phage infection dynamics overtime. Next, we test how antibiotic treatment changes phage-bacteria interactions, both *in vitro* and *in vivo*. Finally, we compare phage-antibiotic synergy across different, mechanistically distinct, antibiotic stressors to begin unveiling the molecular basis of observed phage-antibiotic synergies. Overall, our results suggest that the interactions between individual temperate phage-bacteria pairs are highly context specific, can range from symbiotic to antagonistic depending on environmental conditions, and that temperate phage can mediate how microbiomes respond to perturbations, like antibiotic treatment.

## Methods

### Bee collection and Dissection

To isolate bacteria and phage, foraging honey bees were collected by hand using an insect collection net. All bees were collected near the Harry H. Laidlaw Jr. Honey Bee Research Facility on the University of California Davis campus. After collection, live bees were transported back to the lab and incubated at 4°C for 5-10 mins (enough time for the bees to become inactive). The mid-hind gut region of bees was then dissected in sterile PBS using flame sterilized forceps. After dissection, individual (bacterial isolation) or pooled (viral isolation) bee guts were placed in sterile 1.7mL microcentrifuge tubes on wet ice pending further processing.

### Bacterial Isolation and Cultivation

To isolate bacteria, the mid-hindgut region of individual bees was macerated with 500 µL of sterile PBS using a sterile plastic pestle. The resulting bee gut homogenate was then diluted in 10-fold series to a final concentration of 1×10^-5^. A total of 20 µL of each dilution was then plated on solid MRS agar with 2% fructose (MRS-2F) and 100 mg/L cycloheximide. Plates were incubated anaerobically at 37°C for 72 hours. After incubation, well isolated colonies were picked and streaked for isolation using the same incubation conditions.

To prepare freezer stocks, subcultured bacterial isolates were first grown overnight in liquid MRS-2F. Those overnight cultures were then diluted to an OD_600_ of 0.2 and combined 1:1 with a 30% glycerol solution, with 2 mL aliquots frozen at -70°C for future use. For routine growth, 100 µL of these 0.1 OD_600_ bacterial freezer stocks were inoculated into 3mL of 1x or 2x MRS-2F broth and incubated at 37°C and 5% CO_2_.

### Phage Isolation and Cultivation

Phage isolation and propagation took place using protocols adapted from the Sea-Phages program (https://discoveryguide.seaphages.org/). Between 15 and 30 whole bee guts were pooled into a single 1.7mL microcentrifuge tube containing 500 µL SM buffer (100mM NaCl, 10mM MgSO_4_, 0.01% Gelatin, 50mM Tris-HCL). Pooled bee guts were then macerated using a sterile plastic pestle. Next, this homogenate was transferred to a 50mL conical tube and combined with an additional 9.5mL SM buffer and approximately 5 sterile glass plating beads. Tubes were then incubated horizontally at 4°C for 3 hours at 150 rpm. After incubation, tubes were centrifuged at 3,000 x g for 10 min. The resulting supernatant was then strained using a 0.22 um PES syringe filter. To enrich for phage, 500 µL of an overnight bacterial culture grown in 2x MRS-2F broth was mixed 1:1 with filtered bee gut homogenate and incubated overnight at 37°C 5% CO_2_. Enrichment cultures were then centrifuged at 3,000g for 10 min and filtered using a 0.22 um PES filter. Finally, soft agar overlays were performed using the phage fraction of enrichment cultures by combining 200 µl of a filtered phage enrichment and 100 µl of a fresh overnight culture of the corresponding host bacterium to a tube containing 4 mL of molten MRS-2F soft (0.75% w/v) agar. Soft agar overlay plates were incubated anaerobically at 37°C for 48 hours before checking for the presence of phage plaques.

To isolate phage, well-isolated individual plaques were picked using a flame sterilized agar-plug cutting tool. This agar plug was then transferred to a 1.7 ml tube, combined with 500 µl of SM buffer, and broken up using a sterile plastic pestle. This plug-SM buffer homogenate was then incubated for 3 hours at room temperature at 250 rpm before being centrifuged at 3,000 x g for 10 min. The supernatant was then filtered with a 0.22 um PES filter. This filtrate was then serially diluted out to 10^-5^ and used to perform another round of soft agar overlays. This process was then repeated for 3 total serial passages of individual plaques.

To generate high concentrations of phage for DNA extraction and experimental work, we performed polyethylene glycol (PEG) precipitations of phage lysates generated from webbed plates. First, web plates were generated by performing soft agar overlays (as described above) using 100 µl of a 0.1 OD_600_ fresh bacterial culture and ∼3×10^4^ PFUs. Phage were then extracted by incubating webbed plates with 4mL of SM buffer overnight at 4°C. After incubation, the SM buffer from each plate was filter sterilized using a 0.22 um PES filter and titered via soft agar overlay. To precipitate phage, this high-titer phage lysate was combined 4:1 with a 5x concentration PEG precipitation solution containing 200 g/L PEG8000 and 2.5M NaCl. This solution was then mixed briefly by inverting, incubated overnight at 4°C, and then centrifuged at 20,000 x g for 1 hour at room temperature. The supernatant was then discarded and the pellet resuspended in 1/10^th^ of the original volume of the high-titer phage lysate.

### DNA extraction and Phage genome sequencing

To extract DNA for phage genome sequencing, we followed the protocol used in Sbardellati and Vannette^36^. Briefly, 200 µL of precipitated phage was treated with DNase (20 µL DNase, 20 µL DNase buffer) and incubated at 37°C for 90 mins. Reactions were terminated by adding 20 µL of EDTA and incubating at 65°C for 10 min. We then extracted DNA using a DNeasy PowerSoil Pro kit (Qiagen, Hilden, Germany) following the manufacturer’s instructions with the optional Qiagen Vortex Adapter 15 min bead beating step.

Extracted DNA was submitted to the University of California Davis Genome Center for library prep and paired end 150 bp shotgun metagenomic sequencing. Library prep was performed using a Kapa Hyper prep kit (Kapa Biosystems-Roche, Basel, Switzerland) with Illumina TruSeq adapters (Illumina, San Diego, CA) and a target insert size of 350 bp. Sequencing took place on an Illumina NovaSeq.

Raw Illumina sequencing library was first randomly subsampled to 50,000,000 paired end reads using seqtk^41^. Next, the TrimGalore^42^ was used to remove polyG tails and Trimmomatic^43^ was used to remove adapters and to quality filter raw reads. Cleaned sequencing reads were then assembled using Megahit^44^ and phage predicted with VIBRANT^45^. Our predicted phage genome was then quality checked and trimmed with checkV^46^. Phage taxonomy was predicted using geNomad ^47^. Phage genes were predicted using Pharokka^48^ and genome plot generated using Phold^49,50^.

### Plate reader and growth curve assays

To measure the effects of carbon substrate, phage, or antibiotics on the ability of Amel_191 to grow, we performed a series of growth assays using 96-well microtiter plates. For carbon utilization experiments, a 2x minimal MRS media was made in house by UC Davis Biological Media Services using the recipe provided by Zheng et al.^31^ (2g peptone, 1g yeast extract, 1g polysorbate 80, 0.4g NH_4_Cl, 0.2g MgSO_4_•7H_2_O, 0.07g MnCl_2_•4H2O, 2g KH_2_PO_4_, 0.4g L-cysteine hydrochloride, 0.1g pyridoxine hydrochloride, 0.5g pantothenic acid, 0.1g inositol, 0.01g aminobenzoic acid, and 0.02g adenine). We then combined this media 1:1 with a sterile 2% solution of either glucose, xyloglucan (Megazyme, Bray Business Park, Southern Cross Rd, Bray Co., Wicklow, A98 YV29, Ireland), or arabinogalactan (Vital Nutrients, 45 Kenneth Dooley Drive, Middletown, CT, 06457, USA). Alternatively, we added a pollen extract (50mg/mL) prepared using sterile pollen according to the methods outlined in Kešnerová et al.^30^. For antibiotic-phage synergy experiments, stock antibiotic solutions were prepared by dissolving powdered ampicillin (Thermo Fisher, 168 Third Ave, Waltham, MA, 02451, USA), tetracycline (Millipore Sigma, 400 Summit Drive, Burlington, MA, 01803, USA), or ciprofloxacin (Thomas Scientific, 1654 High Hill Rd, Swedesboro, NJ, 08085, USA) in HPLC grade methanol to a final concentration of 1000 mg/L. These stock concentrations were then diluted to a range of different concentrations such that 100 µL could be combined with 5.9 mL of MRS-2F to generate our target final concentration.

To generate growth curves, we first added 200 µL of different MRS based media to the wells of a 96 well plate. Next, fresh overnight cultures of *Bifidobacterium* were pelleted, washed once with PBS, resuspended in fresh PBS, and then adjusted to a final density of 0.1 OD_600_. These washed bacterial cells were then either combined with an equal volume of phage dilutions or a SM buffer negative control and then incubated for 15 mins. Wells were then inoculated by adding 50 µL of *Bifidobacterium*-phage solutions. After inoculation, carbon utilization plates were placed in an anaerobic environment and incubated at 37°C. Growth, quantified as OD_600_, was measured once at 48 hrs using an automatic spectrophotometer plate reader (Biotek HTX, Agilent Technologies, 5301 Stevens Creek Blvd, Santa Clara, CA, 95051, USA). Plates testing antibiotic synergy were placed in the same plate reader and incubated at 32°C for 120hrs. Every 15 mins, plates were shaken for 15 sec (6-mm diameter at 6 Hz) and OD_600_ was measured.

### Bee rearing and laboratory cage setup

Bees for gnotobiotic experiments were sourced from hives maintained by the USDA ARS Pollinator Health Unit in Davis, CA and reared using protocols adapted from previous publications^31,32^. To ensure that gnotobiotic bees would emerge at roughly the same time, queens from 3-4 separate colonies were caged and permitted to lay eggs for 48 hours on frames containing drawn wax comb, but without developing brood or stored food. After 16 days, these brood frames, now containing capped pupae, were removed from colonies and brought into the lab. Pupae with pigmented eyes, but lacking movement, were then transferred to the wells of sterile 48 well plates and incubated at 34.5°C and 75% relative humidity. Plates of developing pupae were checked twice daily for the presence of emerged adult bees. Each day, adults originating from the same source colonies were placed into a sterile cup cage and provided *ab libitum* access to a sterile 0.5M sucrose solution, similar to what was described by Zheng et al.^31^.

For laboratory cage setups, adult bees which had emerged over a 48 hr time-period and originated from the same source colonies were pooled together. Pools of adult bees were then divided into sets of 20-30 bees and randomly distributed into new sterile cup cages and incubated at 34.5°C and 75% relative humidity. At this point, 1 bee from each cage was randomly sampled and used to confirm microbiota deplete status via qPCR. For *Bifidobacterium*, primers targeting the 16S gene were adopted from Kesnerova et al.^30^. For phage, custom primers targeting the tail tape measure gene of phage Isabelle were designed in house using NCBI’s primer blast tool (Forward: CAGCCAGGCAGGCAAAGTAGCATCCTTCCCAGGTCCCTTT; Reverse: AAGCTCGGCATCCACTCTCCATTTGGGAAAGGCTGCGATTG). All qPCR reactions were performed following the method described in Christensen et al.^51^. Each reaction contained: 5 µL SSO Advanced Universal SYBR Supermix 41 (Biorad, Hercules, CA, USA), 0.5 µL of forward primer (5 µM), 0.5 µL reverse primers (5 µM), 3 µl molecular grade water, and 1 µl of template DNA (diluted 1:10 in molecular grade water). Reactions were performed in triplicate for each sample.

### Inoculation of bee cages

Bacterial inocula were prepared by centrifuging thawed 15% glycerol stocks (3,000 x g, 15 min, 4°C), discarding the supernatant, and resuspending the pellet in PBS to a final OD₆₀₀ of 1.0 (∼1.5 × 10⁶ cells/µL). Phage inocula were prepared via PEG precipitation and SM buffer resuspension of high-titer lysates (as described above) and adjusted to ∼5 × 10⁵ PFU/µL. Microbial treatments were prepared identically across both experiments.

Established microcolonies were fed sterile 0.5M sucrose for their first 24 hrs. Bees were then starved for 4 hours before being provided 24 hour *ad libitum* access to microbial treatments. Treatments differed according to experiment (**Fig. S6 and S9**). For the first in-vivo experiment, bees were provided with a solution containing 500 µL 50% sucrose (w/w) with either: 1) 250 µL PBS and 250 µL SM buffer, 2) 250 µL *Bifidobacterium* and 250 µL SM buffer, 3) 250 µL *Bifidobacterium* and 250 µL phage, or 4) 250 µL PBS and 250 µL phage. In our second experiment, bees were provided a solution containing 500 µL 50% sucrose (w/w) with either: 1) 250 µL *Bifidobacterium* and 250 µL SM buffer, or 2) 250 µL *Bifidobacterium* and 250 µL phage. Following inoculation, all microcolonies received *ad libitum* access to 0.5 M sucrose (replaced every 48 hrs) and sterile pollen (replaced every 72 hrs). In the second experiment, beginning 72 hrs after the switch to standard diet, sucrose solutions were replaced with solutions of 0.5 M solutions containing varying dosages of tetracycline for a 48 hr treatment window. Tetracycline-sucrose solutions were replaced daily.

### Honey bee sampling and microbiological quantifications

For the first in-vivo experiment investigating bacteria and phage colonization and interactions within the bee gut, 3 bees were sampled from each cage at 24, 48, 120, 192, and 264 hrs post pollen introduction. For our second in-vivo experiment, which examined how antibiotics alter bacteria-phage interactions, 3 bees were sampled from each microcolony at 3 different time points: 24 hrs pre antibiotic treatment, immediately following antibiotic treatment, and 72 hrs post antibiotic treatment.

Bees were collected from cup cages using sterile forceps, placed into 15 mL conical tubes, and transferred back to the lab on ice where they were stored at −20°C pending further processing. Prior to dissection, bees were thawed on wet ice. The mid-hindgut was dissected in sterile PBS using flame-sterilized forceps. Dissected gut material was then weighed and homogenized (4.5 m/s, 30 s) in 500 µL sterile molecular-grade water using a Benchmark Scientific BeadBlaster 24 Microtube homogenizer (Benchmark Scientific, 2600 Mian Street Extensions, Sayreville, NJ, 08872, USA). For DNA extractions, 200 µL of homogenate was stored at −70°C until further processing. DNA was extracted from bee gut homogenate using a DNeasy PowerSoil Pro kit (Qiagen, Hilden, Germany) following the manufacturer’s instructions with an optional Qiagen Vortex Adapter 10 min bead beating step. Bacterial and phage loads were determined from extracted DNA via qPCR using organism specific primers, as described above.

### Statistical and computational analysis

All statistical analyses were conducted in R^52^ v4.2.3. One-way ANOVA and TukeyHSD posthoc tests were used to test how *Bifidobacterium* growth differed across carbon substrates and how *Bifidobacterium* abundance differed *in vivo*. T-tests were used to test how phage impacted bacterial growth *in vitro*. Gene-sharing networks were generated outside of R using the program vConTACT2^53^. Genome alignment plots were constructed using clinker^54^.

## Results

### Bees House Complex Carbohydrate Degrading *Bifidobacterium*

To begin studying phage-bacteria interactions in the honey bee gut, we first sought to isolate suitable bacterial hosts. To this end, we plated honey bee gut homogenate on MRS agar supplemented with 2% fructose. This yielded a collection of different bacterial isolates which we presumptively identified as strains of *Bifidobacterium* and *Lactobacillus* based on cell morphology. From these, we chose a single isolate, Amel_191, for further characterization.

To further identify Amel_191, we performed Sanger sequencing targeting the full-length 16S rRNA gene (**Table S1**) and constructed a phylogeny comparing this gene to those found in other *Bifidobacterium* spp. (**Fig. 1A**). This presumptively identified Amel_191 as a strain of *B. polysaccharolyticum*, a recently described taxa of bee-associated *Bifidobacterium* related to *B. asteroides*^55^.

**Figure 1:**
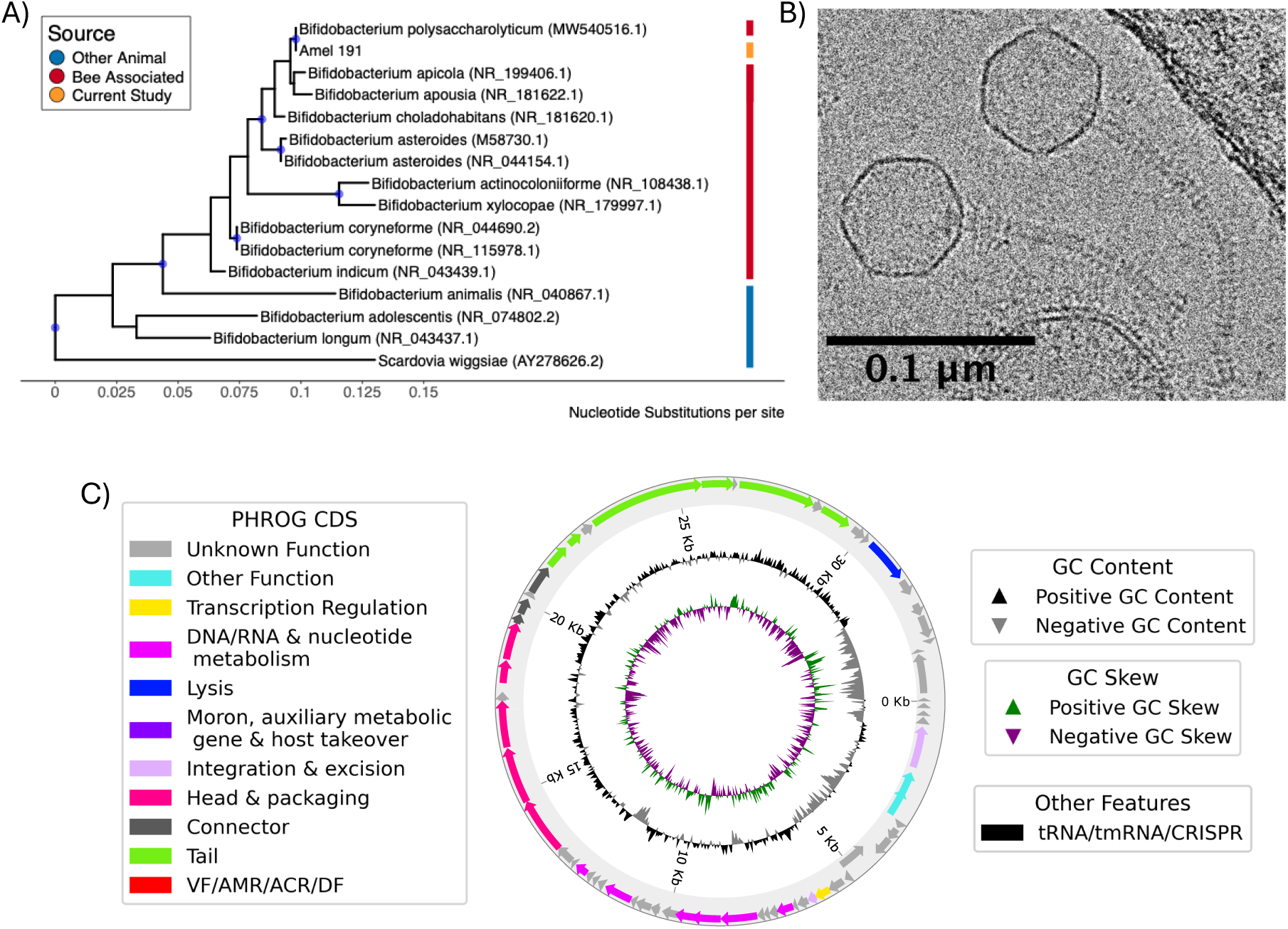
Description of bacteria and phage isolated in the current study. A) Phylogenetic tree describing the relatedness of Amel_191 to other animal associated *Bifidobacterium*. Color ribbon indicates host animal type. Nodes with bootstrap support >= 75% are colored with blue points. Tree was built using full length 16S sequences. Scale indicates the number of expected nucleotide substitutions per site. B) Cryo-electron micrograph describing the morphology of phage Isabelle. Scale bar represents 100 nm. C) Full genome sequence of phage Isabelle visualized by Phold and Circos. Outer ring contains predicted genes colored by function. Arrows correspond to individual genes. Middle ring represents GC content and inner most ring represents GC Skew.

Bee-associated *Bifidobacterium* are believed to play an important role in allowing bees to derive nutrition from the structural carbohydrates present in pollen cell walls^31^. To test if the *Bifidobacterium* we isolated can perform this metabolic action, Amel_191 was grown in modified MRS media containing either arabinogalactan, xyloglucan hemicellulose, or a water-soluble pollen extract as carbon sources (**Fig. S1**). The results of this experiment demonstrated that Amel_191 grew in minimal media containing pollen, or pollen associated polysaccharides, as the main carbon sources (Arabino vs No sugar p < 0.001, Xylo vs No sugar p > 0.05, Pollen vs No sugar > 0.001), highlighting the potential role of this bacterium in bee nutrition.

### Temperate Phage Infection Modulates *Bifidobacterium* Growth on Complex Carbohydrates

To isolate a phage capable of infecting Amel_191, phage enrichments and soft agar overlays using Amel_191 and the phage fraction of adult honey bee gut homogenate were performed. Following this approach, a single phage was isolated which we have named Isabelle.

Phage Isabelle forms small (∼1.8 mm), turbid, punctiform plaques when grown anaerobically on MRS soft agar overlays (**Fig. S2**). Cryo-Electron Microscopy revealed Isabelle to be a tailed phage, with a head of ∼55 nm and a tail of 100-150 nm (**Fig. 1B**). Following short read sequencing and genomic assembly, Isabelle was found to possess a circular genome of 34,782 bp (**Fig. 1C**), which is predicted to encode 63 coding sequences. Similar to what has been described for other phage, the genome of Isabelle appears to be organized into discrete modules with specific functions. To better understand the placement of the isolated phage within the greater landscape of bee-associated phage, we determined its taxonomy using geNomad^47^ and visualized gene-sharing between it and phage previously described in bee gut metagenomes (**Fig. S3**). This analysis identified phage Isabelle as a Caudoviricetes and placed it amongst other *Bifidobacterium* phage found in bees, particularly those described in Bonilla-Rosso et al.^34^ and our own previous work^36^ (**Fig. S4**). Lastly, we note that Isabelle is predicted to encode an integrase gene which, when combined with the observed turbid plaque morphology, suggests a temperate lifestyle.

Equipped with a phage-bacteria pair, we next wanted to test how phage infection might influence gut-microbiome function. To address this question, Amel_191 was cultured in modified MRS broth supplemented with different carbon sources, but this time included phage Isabelle at a range of different concentrations (**Fig. S5**). This experiment revealed a substrate and phage-dosage dependent pattern. Overall, high concentrations of phage negatively impacted the growth of Amel_191 on glucose (t_4_ = 13.762, p < 0.001), pollen ( t_4_ = 30.679, p < 0.001), and arabinogalactan (t_4_ = 16.282, p < 0.001). However, this same phage treatment had no impact when bacteria were growing on xyloglucan (t_4_ = 0.12263, p > 0.05). Similarly, at lower dosages, phage had little to no impact on bacterial growth. In fact, low phage dosages appeared to moderately increase bacterial growth when Amel_191 was grown on either arabinogalactan or xyloglucan (**Fig. S5**), though these differences were not significant.

### Phage Isabelle Improves *Bifidobacterium* Colonization of Bee guts

Prompted by the observation that phage effects on Amel_191 change in a substrate-dependent manner, we next sought to characterize phage-bacteria interactions *in vivo*. To accomplish this, microbiota-depleted bees were inoculated with phage and bacteria according to a 2×2 factorial design (**Fig. S6**) and bacterial and phage abundance monitored over time using qPCR (**Fig. 2A**).

**Figure 2:**
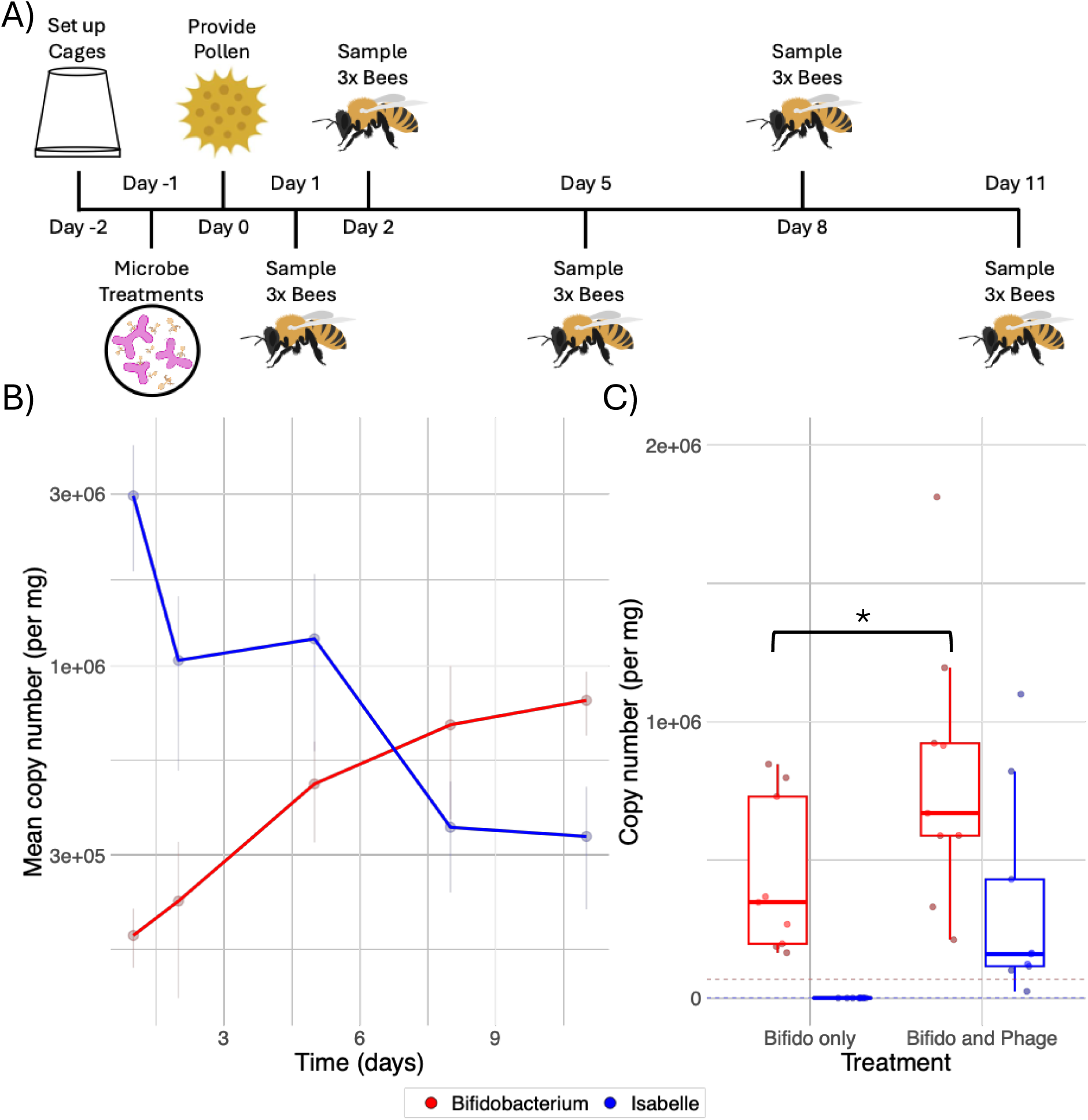
Phage abundance declines over time, but improves bacterial monocolonization of the honey bee gut. A) Timeline representing sampling scheme of in vivo experiment. Note, “Day 0” is defined by the switch of bees to their standard diet. B) Line plot describing how *Bifidobacterium* abundance (red) and phage abundance (blue) changes overtime. Points represent averages +/- standard error. Time is represented along the x-axis, while microbe copy number is shown along the y-axis. C) Boxplots visualizing the abundance of *Bifidobacterium* (red) and phage Isabelle (blue) at the end of the experiment. Treatment (Bifidobacterium +/- phage) on the x-axis. Copy number is along the y-axis.

Overall, we observed that *Bifidobacterium* density increased steadily throughout the experiment, indicating successful colonization of bee guts, while phage abundance decreased over time (**Fig. 2B**, **Fig. S7**). Interestingly, bees receiving both Amel_191 and phage Isabelle exhibited significantly higher densities of *Bifidobacterium* compared to those receiving bacteria alone (Figure 2C, Bifido vs Bifido and Phage p < 0.05), suggesting that phage presence enhanced bacterial colonization efficiency or population growth. However, this increase is modest and relatively variable, with a few individuals likely driving the trend. We screened for phage or bacterial contamination and detected low bacterial numbers in our phage only treatment (**Fig. S7**). Because this contamination was only present at the final time point and in a single bee cage, we retained these samples in our analysis.

To further investigate the mechanism behind why *Bifidobacterium* abundance did not decrease following phage treatment, the virus-to-microbe ratio (VMR) of phage Isabelle and *Bifidobacterium* was examined *in vivo* (**Fig. 2B** and **S8**). Results indicated that, at early timepoints, phage outnumber their bacterial hosts approximately 10 to 1, but that this ratio decreased rapidly over time, dropping below 1 by approximately day 6. This suggests that the infection strategy employed by Isabelle changed throughout the course of the experiment, likely shifting from lytic at early time points to lysogenic at later time points.

### Phage Killing of *Bifidobacterium* is Enhanced by Antibiotic Stress

To investigate if antibiotics altered phage-bacteria interactions, we grew Amel_191 in the presence of varying concentrations and combinations of phage Isabelle and tetracycline, a protein synthesis inhibiting antibiotic. We chose tetracycline for this work because of its common use in commercial apiaries. This experiment revealed a synergy between phage Isabelle and tetracycline (**Fig 3A**). For example, while sublethal tetracycline treatment (0.1 ug/mL) alone produced little to no change in bacterial growth, if the same antibiotic dosage was paired with even a low titer of phage (>10 pfu) bacterial growth was strongly suppressed, reaching ∼50% of untreated controls by 192 hours (**Fig 3B**).

**Figure 3:**
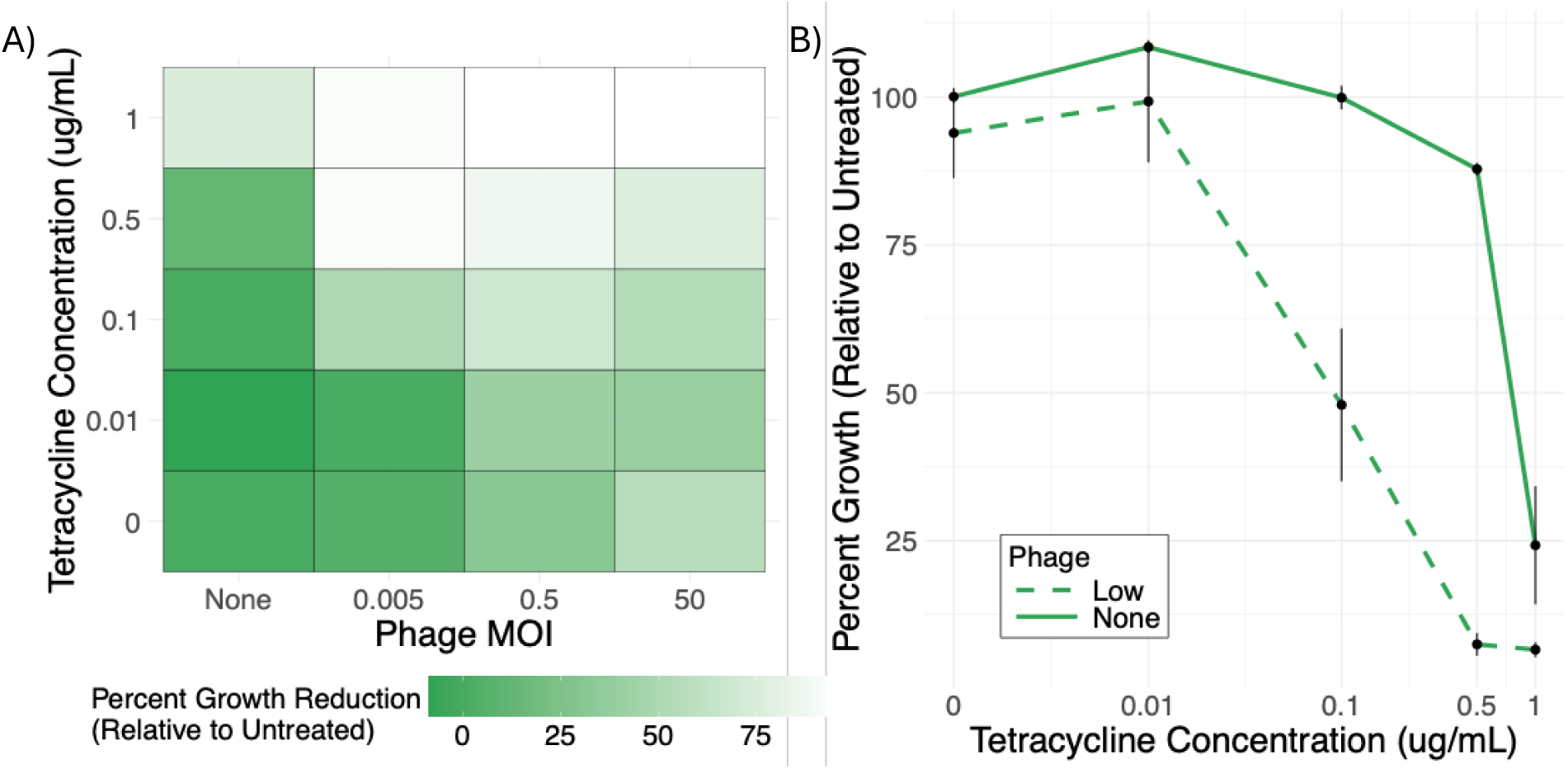
Phage Isabelle acts synergistically with tetracycline. A) Checkerboard assay presenting phage Isabelle-tetracycline synergy across a range of different phage (x-axis) and antibiotic (y-axis) concentrations. Color represents bacterial growth, expressed as percent growth relative to untreated bacteria (grown without antibiotics or phage). B) Line plot providing a more detailed look of data seen in panel A. Increasing tetracycline concertation is represented along the x-axis. Percent growth, relative to untreated bacteria, depicted on y-axis. Points represent averages +/- standard error. Solid lines show Amel_191 growth without phage, while broken lines show growth with a low dosage of phage. Note, the x-axis is log-transformed to improve legibility.

Building on the observation that antibiotic stress alters phage-bacteria interactions *in vitro*, we next hypothesized that phage and antibiotics might also interact synergistically to suppress bacterial growth *in vivo*. To test this, we set up another *in vivo* experiment where microbiota-depleted bees were treated with Amel_191, phage Isabelle, and varying dosages of tetracycline according to a 2×4 factorial design (**Fig. S9**). We then monitored bacterial and phage abundance over time using qPCR (**Fig 4A**). Overall, phage presence significantly impacted the effect of antibiotic treatment on *Bifidobacterium* abundance (**Fig. 4B**). Bees colonized with only Amel_191 showed no significant change in *Bifidobacterium* abundance following tetracycline treatment (p > 0.05), although high concentrations (450 ug/mL) qualitatively reduced bacterial density. In contrast, when bees were inoculated with both Amel_191 and phage Isabelle, *Bifidobacterium* abundance was significantly decreased across all tetracycline dosages, relative to no antibiotic controls (**Fig. 4B**; all p < 0.05). However, despite this apparent synergy, we observed little to no corresponding change in VMR (**Fig. 4C**), suggesting that phage induction by antibiotic treatment was not necessarily responsible for the observed phenotype.

**Figure 4:**
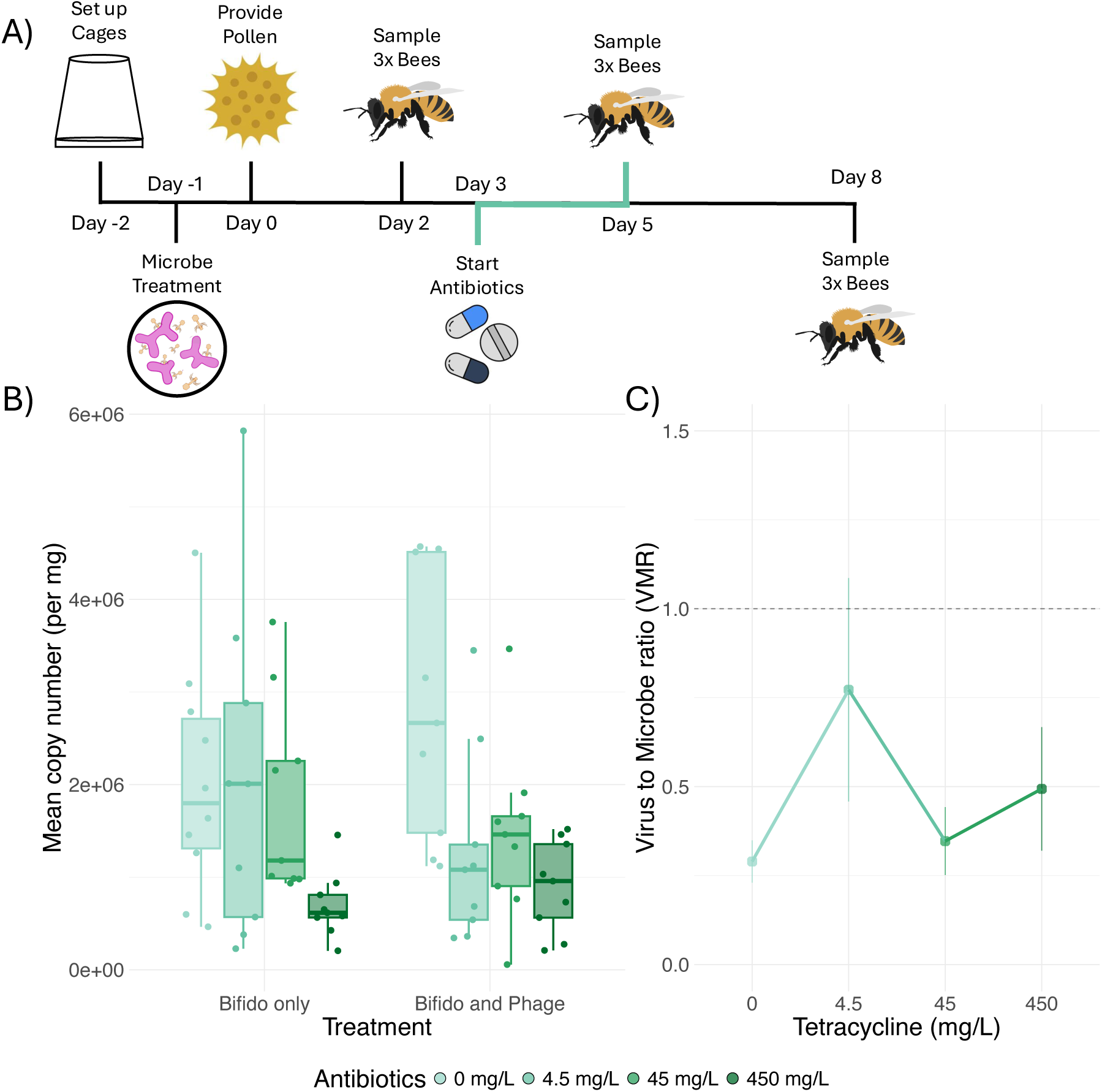
Phage-antibiotic synergy occurs in *in vivo*. A) Timeline representing sampling scheme of second *in vivo* experiment. Note, “Day 0” is defined by the switch of bees to their standard diet. B) Boxplots describing the abundance of *Bifidobacterium* in bee guts at the end of the experiment. Treatment group (*Bifidobacterium* +/- phage Isabelle) represented along x-axis. Bacterial abundance, as measured by qPCR, shown on y-axis. Color represents increasing tetracycline concentration, with darker colors indicating high antibiotic concentrations. C) Line plot depicting how the virus-to-microbe ratio (VMR) in bees treated with both *Bifidobacterium* and phage differed by tetracycline dosage. Antibiotic concentration is on the x-axis, VMR is on the y-axis. A dotted line is drawn at VMR = 1, meaning a equal number of phage and host bacterium. As before, color represents tetracycline concentration.

### Mechanism of Phage-Antibiotic Synergy Differs by Antibiotic

Intrigued by the observation that tetracycline alters the way phage-bacteria pair interact, we next examined if phage-bacteria interactions were similarly modulated in the presence of other antibiotics with distinct modes of action. When challenged with tetracycline, low and medium phage titers produced growth curves which simply plateaued at lower densities than either tetracycline or phage alone (**Fig. S10**). In contrast, combining the cell-wall damaging beta-lactam ampicillin with medium to high phage dosages lead to steep decline and near clearance of bacteria from culture, indicative of active bacterial lysis by phage-antibiotic synergy (**Fig. S11**). Finally, little to no synergy was observed with ciprofloxacin, a DNA damaging antibiotic. In fact, this antibiotic appeared to mitigate the effect of high phage dosages on bacterial growth, indicating an antagonistic relationship with phage Isabelle (**Fig. S12**). Overall, the results of these experiments show that multiple antibiotics can act synergistically with the phage described here, but that the nature of these interactions can differ substantially by antibiotic.

## Discussion

This research used the honey bee gut as a model to showcase the dynamic nature of phage-bacteria interactions in the gut, and highlights the importance of temperate phage in shaping gut microbial ecology, especially in response to antibiotic exposure. First, we showed that temperate phage infection strategy likely changes throughout the course of bacterial gut colonization, moving from lytic replication to lysogeny. Second, we demonstrate that infection by temperate phage can improve the ability of beneficial gut bacteria to colonize the gut. Finally, we use *in vitro* and *in vivo* experiments to show the importance of phage in shaping how gut microbiomes respond to perturbation by antibiotic treatment. Overall, this work improves our fundamental understanding of phage in the gut, thereby shedding light on the basic factors shaping gut microbiomes.

The work performed here builds on existing research by providing experimental validation for observations made by previous metagenomic surveys. For example, a recent study of healthy Danish infants, adolescence, and adults used metagenomic sequencing to show that the abundance and diversity of temperate phage in the human gut decreases over time^56^. In agreement with this, our results revealed a steady decrease in the VMR between temperate phage Isabelle and its *Bifidobacterium* host as bees aged. Taken together, these results suggests that the lytic activity of temperate phage decreases as microbial density increases and gut microbiomes mature. Our data also suggest that temperate phage infection can improve bacterial colonization in the gut, adding to a growing body of literature which implicates phage as important mediators of bacterial ecological fitness^57,58^. While previous studies have shown that phage infection can promote traits associated with bacterial persistence, including biofilm production^59,60^ and spore formation^61^, our work is among the first to demonstrate that phage can directly enhance bacterial colonization within the gut. Future work may investigate this further by investigating the mechanisms by which phage promote bacterial gut colonization.

Our results also extend our understanding of how phage-antibiotic synergies occur within the gut. Prior work has shown that beta-lactam treatment reshapes gut viral communities^14^, and that phage communities can shape how gut microbiomes respond to antibiotics^15^. Congruent with this, we found that the presence or absence of a single temperate phage substantially modulated the way that *Bifidobacterium* responded to antibiotic challenge. Critically however, we observed that antibiotic treatment produced little to no change in VMR *in vivo*, indicating that the phage-antibiotic synergy we observed occurred independent of temperate phage induction. This simultaneous synergy without induction aligns with findings from Al-Anany et al.^20,21^ and Fatima et al.^12^ who have extensively studied temperate phage–antibiotic synergy *in vitro* and have demonstrated that these interactions can instead arise through a range of diverse mechanisms, including initial lysogeny formation.

Work with both virulent and temperate phage has shown that phage-antibiotic synergies are often complex and mechanistically distinct across different combinations of bacteria, phage, and antibiotics^11,12,20,21^. In agreement with this, we found that challenging phage-bacteria pairs with different antibiotics produced growth curves characteristic of different forms of synergy. While phage-tetracycline interactions simply suppressed maximum bacterial density, yielding growth curves which plateau at lower densities, phage-ampicillin interactions crashed bacterial density, leading to near-total clearance from culture. In contrast, there were no interactions between phage Isabelle and ciprofloxacin, a DNA damaging antibiotic known to elicit bee the SOS response and to synergize strongly with different temperate phage^12,21^. We interpret these results as showing that an individual phage can synergize with different antibiotics in mechanistically distinct ways. Future work could focus on discerning the specific mechanisms for these synergies and may leverage additional information, such as genomic content, to potentially better predict these interactions.

An important caveat with much of the *in vivo* work performed here is that the bees used here were mono-colonized with individual bacteria and phage. Although this design allowed us to focus on specific interactions between bacteria and phage, our results may differ if other bacteria were present to interact with our *Bifidobacterium* isolate. We predict that competition with other bacteria could enhance the declines in *Bifidobacterium* abundance we observed in response to phage-tetracycline treatment. This would be especially true of bacteria which occupy a similar ecological niche but are resistant to infection by phage Isabelle, such as other strains of *Bifidobacterium*. Recent work by Ndiaye et al.^38^ has highlighted the relevance of these interactions by showing that phage host ranges are relatively narrow in the bee gut microbiome, with phage diversity closely tracking bacterial diversity at the strain level. Alternatively, if bees were colonized with a more complete microbiome, we may have observed off-target effects of phage and antibiotic challenges. Gut microbiomes are highly interconnected, with different bacteria often working together to co-operatively breakdown different substrates^62,63^. Consistent with this idea, previous work in mice has shown that phage killing of individual bacteria can negatively impact taxonomically disparate bacteria^64^, likely because of this interconnectedness. Understanding how these individual phage-bacteria interactions impact larger gut microbiomes will be imperative for understanding the role phage play in shaping gut microbiomes and the biotechnological use of phage.

## Availability of Data and Materials

The raw sequencing data generated in this study is available on NCBI under the Bioproject accession number PRJNA1476334.

Other data and source code are available at: https://github.com/dsbard/Bifido_phage.

## Acknowledgements

We thank Richard Martinez and the UC Davis Harry H. Laidlaw Jr. Honey Bee Research Facility staff, as well as Eliza Litsey and members of the Davis USDA ARS Pollinator Health Unit, for help sampling honey bee colonies. We also thank members of the Vannette, Emerson, and Brown lab for manuscript comments and feedback and the UC Davis Genome Center for DNA sequencing. Mention of trade names or commercial products in this publication is solely for the purpose of providing specific information and does not imply recommendation or endorsement by the U.S. Department of Agriculture. USDA is an equal opportunity provider and employer.

