## Supplemental Figures for "Temperate Phage Shape Honey Bee Gut Microbiome Structure and Response to Antibiotic Treatment"

| Scientific Name | Max Score | Total Score | Query Cover | E value | Per. ident | Acc. Len | Accession |
|---|---|---|---|---|---|---|---|
| *Bifidobacterium polysaccharolyticum* | 2473 | 2473 | 100% | 0 | 99.56 | 1423 | NR_181621.1 |
| *Bifidobacterium apicola* | 2451 | 2451 | 100% | 0 | 99.26 | 1390 | NR_199406.1 |
| *Bifidobacterium choladohabitans* | 2425 | 2425 | 100% | 0 | 98.96 | 1427 | NR_181620.1 |
| *Bifidobacterium apousia* | 2412 | 2412 | 100% | 0 | 98.74 | 1428 | NR_181622.1 |
| *Bifidobacterium asteroides* | 2409 | 2409 | 100% | 0 | 98.74 | 1369 | NR_044154.1 |
| *Bifidobacterium mizhiense* | 2398 | 2398 | 100% | 0 | 98.59 | 1425 | NR_179386.1 |
| *Bifidobacterium indicum* | 2362 | 2362 | 100% | 0 | 98.08 | 1514 | NR_043439.1 |
| *Bifidobacterium coryneforme* | 2353 | 2353 | 100% | 0 | 98 | 1370 | NR_115978.1 |
| *Bifidobacterium favimelis* | 2348 | 2348 | 100% | 0 | 97.92 | 1411 | NR_199430.1 |
| *Bifidobacterium coryneforme* | 2268 | 2268 | 100% | 0 | 96.29 | 1473 | NR_044690.2 |

Table S1: Top 10 NCBI Blast result of aligning the full length 16S rRNA sequence of Amel_191 against NCBI's 16S/ITS database.

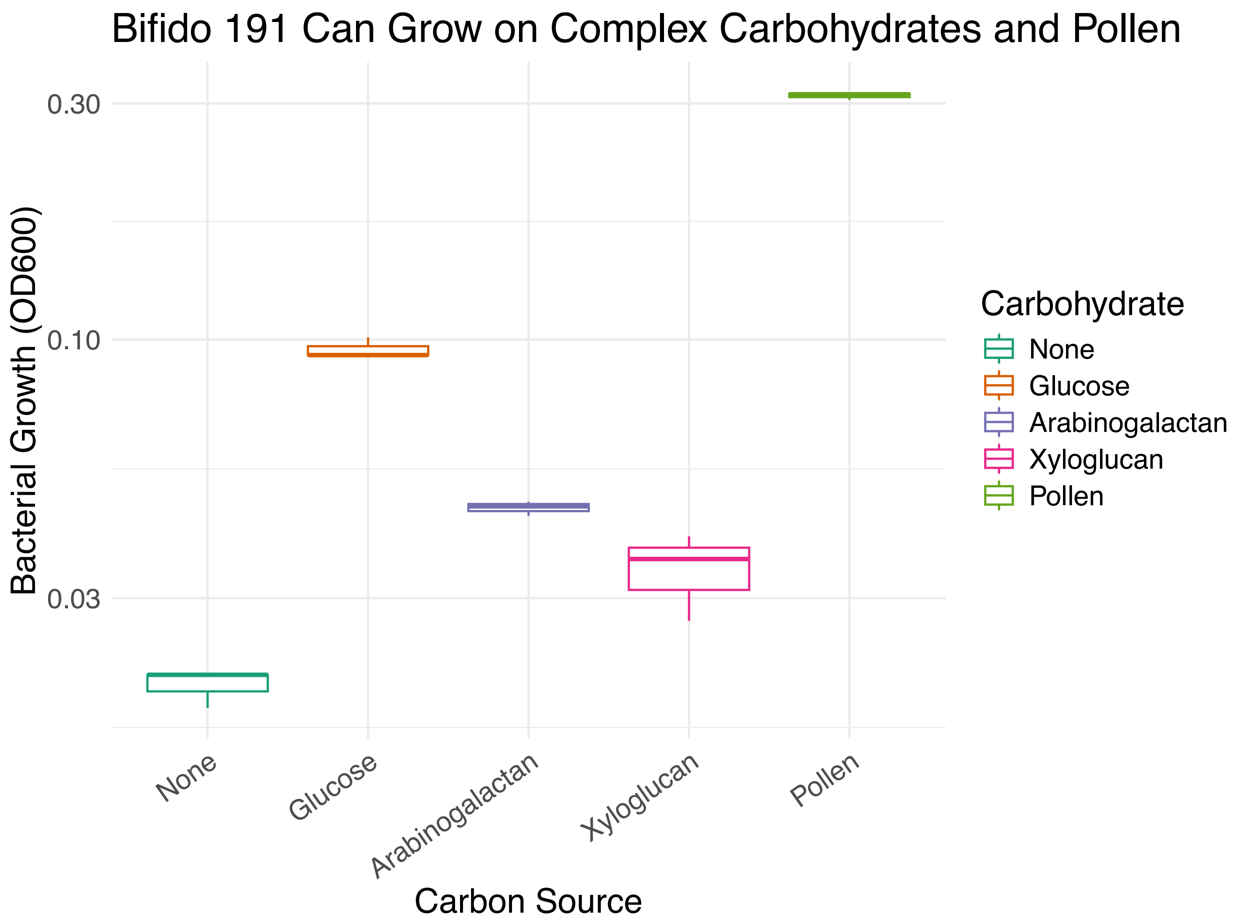


Figure S1: Boxplots describing the growth of Amel_191 in minimal MRS media containing a variety of different carbons sources. Carbon source is on the x-axis, bacterial growth along the y-axis.

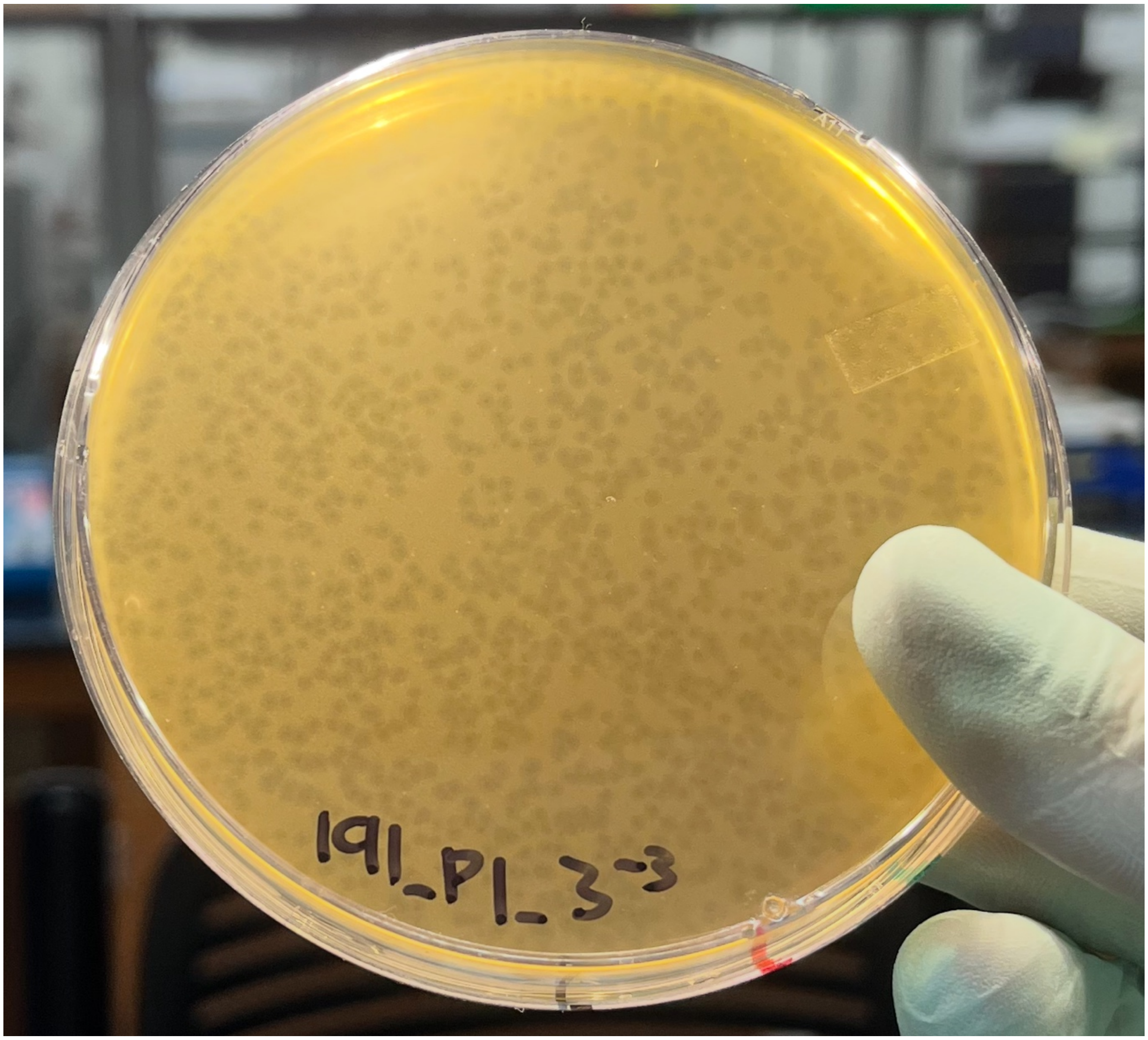


Figure S2: Soft agar overlay showing the plaque morphology of phage Isabelle. Plates were incubated anaerobically at 37C for 48 hrs using Amel_191 as the host.

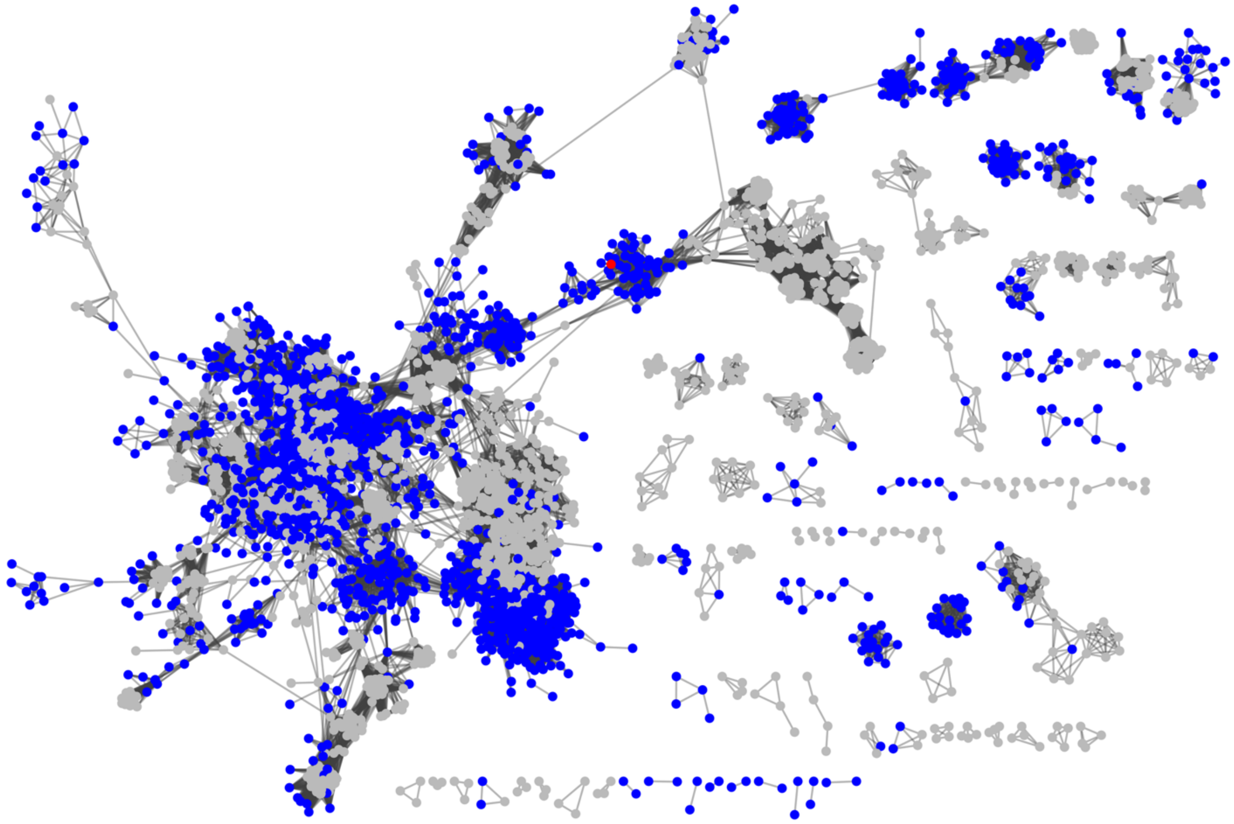

Figure S3: Weighted gene-sharing network of phage Isabelle, reference phage, and all non-singleton vOTUs identified in previous bee phage studies. Individual nodes are phage genomes. Nodes are connected by edges when genomes share genes. Nodes are colored based where phage were identified. Phage Isabelle is highlighted in red. Previously described bee phage are in blue. Reference phage are in grey.

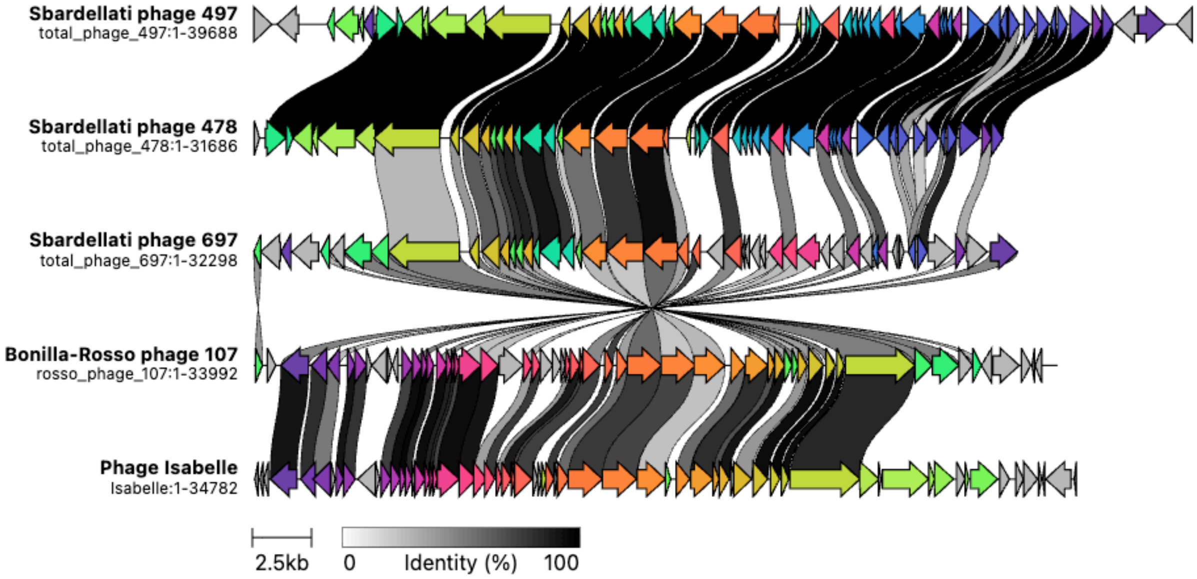


Figure S4: Clinker plot displaying gene-sharing between phage Isabelle and phage which clustered with Isabelle in previous gene-sharing network. Notably, all co-clustering phage were described from honey bee gut material.

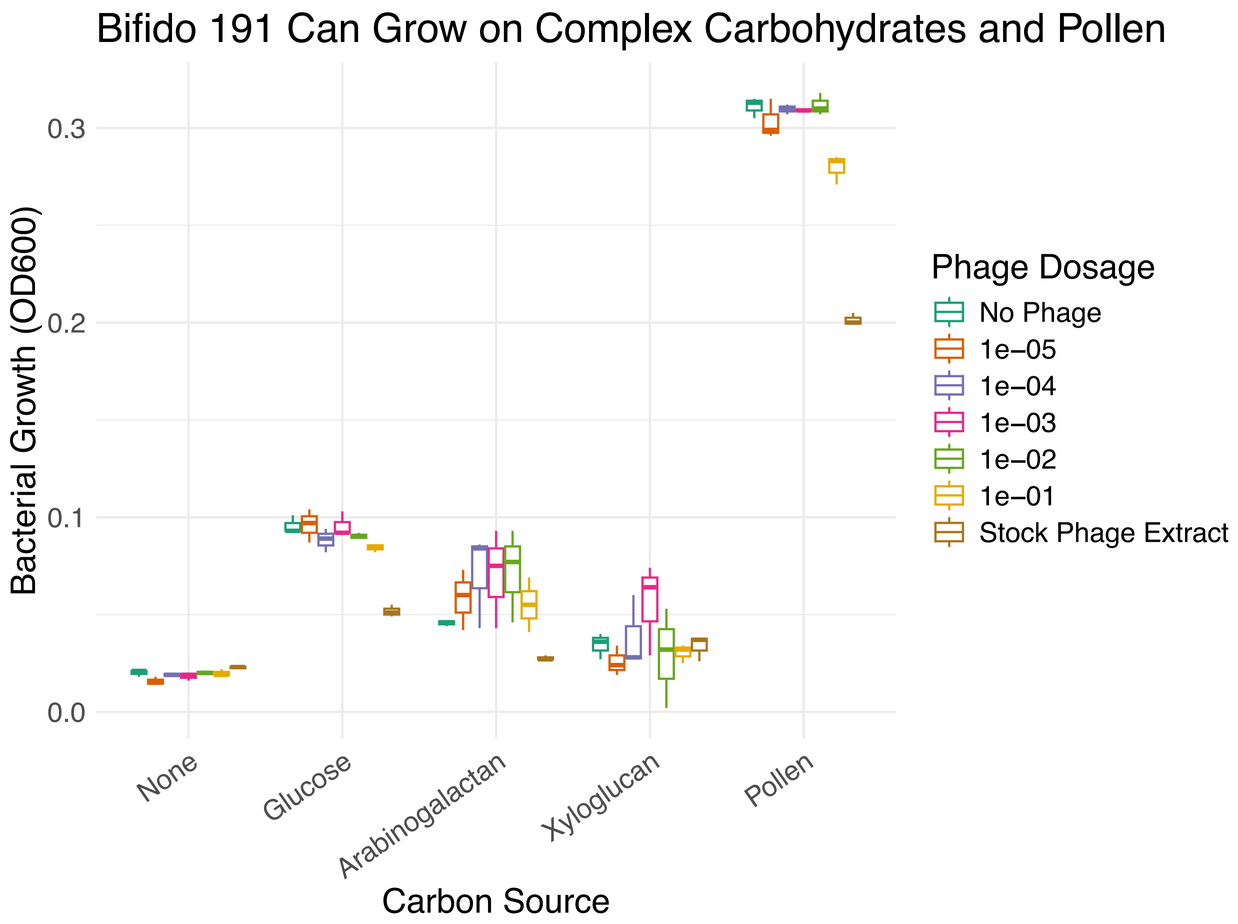


Fig. S5: Boxplots describing how Amel_191 growth on different carbon substrates is affected by different concentrations of phage Isabelle. Bacterial growth (OD600) is represented along the y-axis. Carbon substrate is on the x-axis. Phage dosage is a 10-fold dilution series with 1 being our stock phage extract and 0 being a SM buffer control.

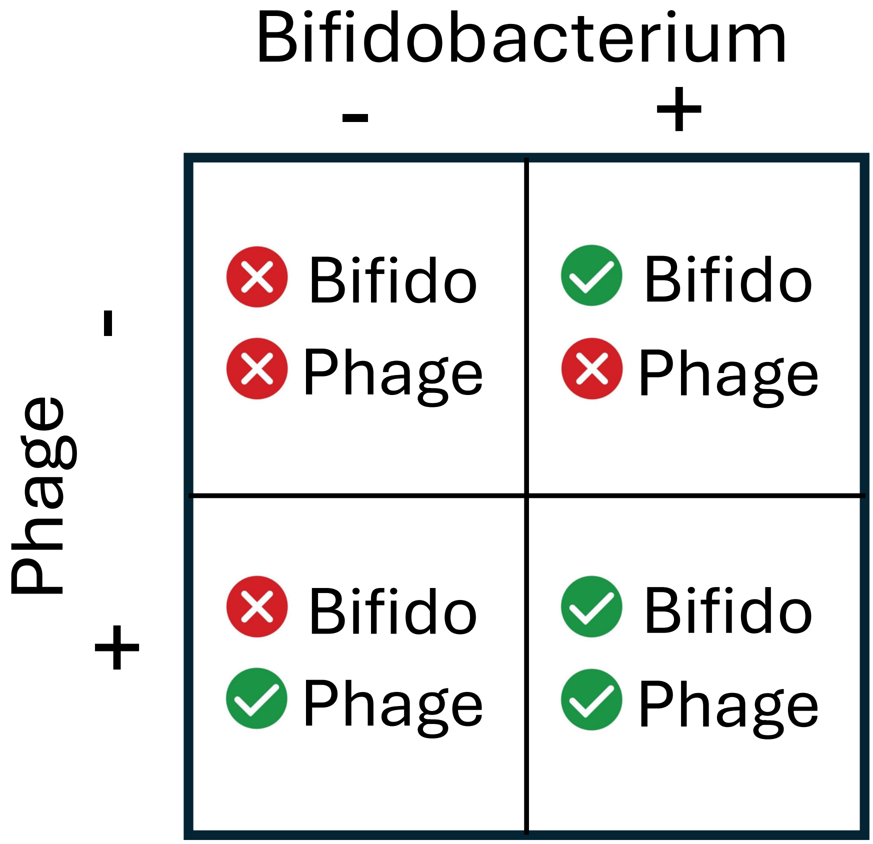


Fig. S6: Schematic representing the 2x2 factorial design used in the first in vivo experiment.

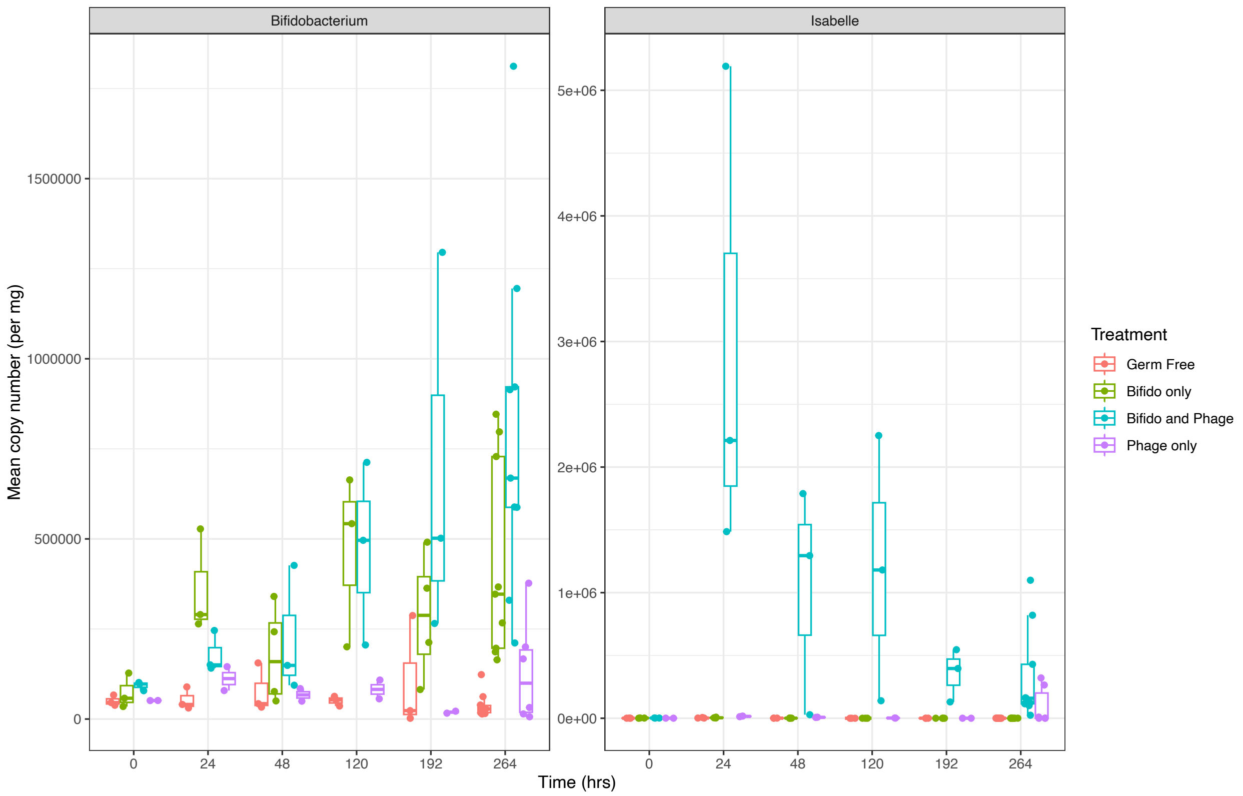


Fig S7: Boxplots describing change in bacterial and phage abundance over time. *Bifidobacterium* abundance is on the left, phage abundance on the right. X-axis is time (in hours) and y-axis is density (copy number per mg gut material). Color denotes treatment type. Note, in bees which received only phage treatments, there is a low abundance of both *Bifidobacterium* and phage at the final sampling time. However, this was only true of animals sampled from one bee cage.

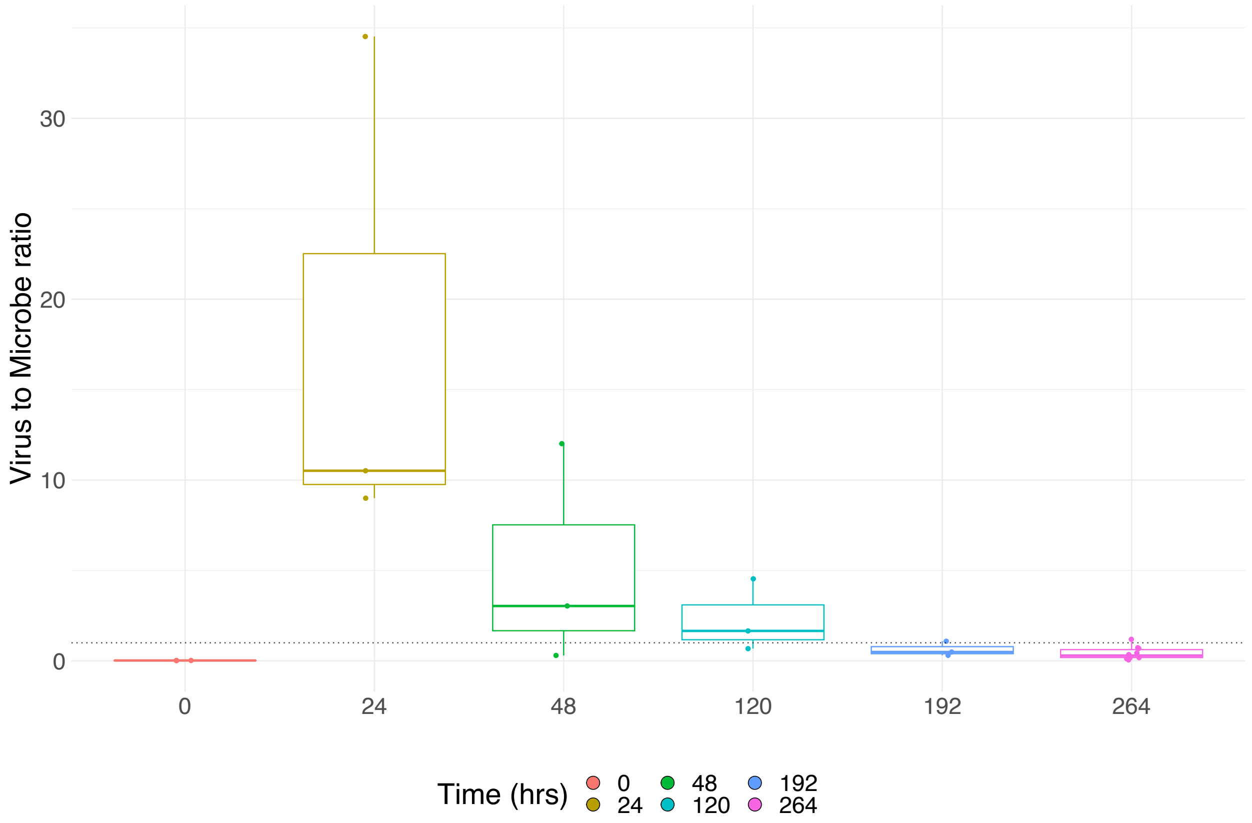


Fig. S8: Boxplots showing the change in virus-to-microbe (VMR) ratio over time in bees treated with both *Bifidobacterium* and phage Isabelle. VMR is on y-axis. Time (in hours) is represented along x-axis.

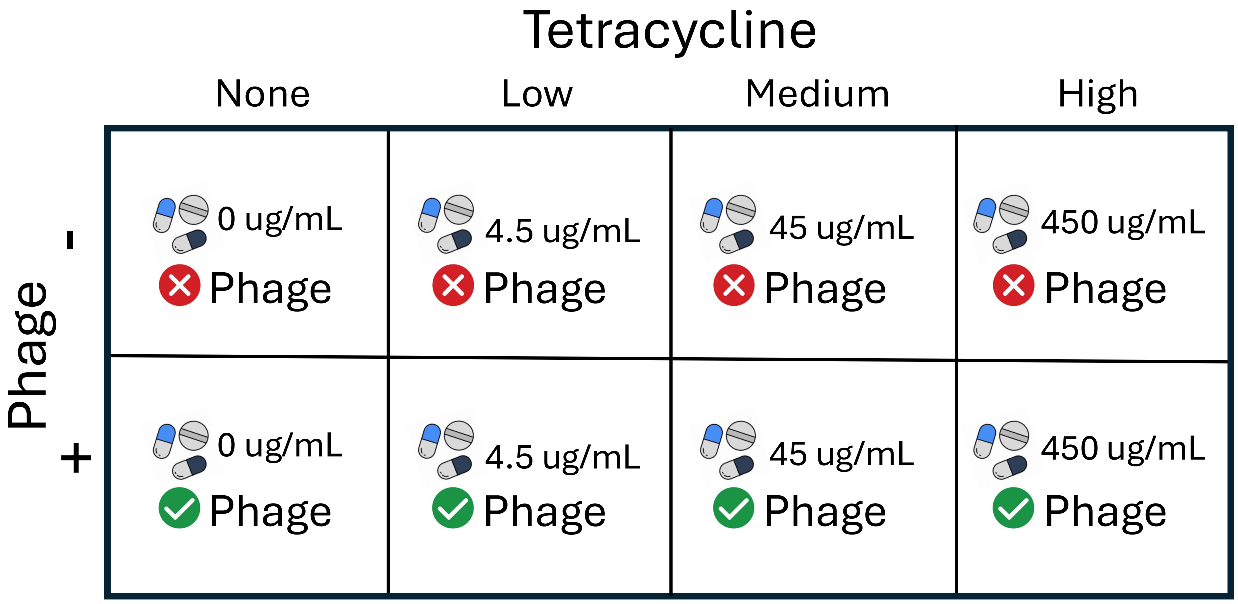


Figure S9: Schematic describing the 2x4 factorial design used in our second in vivo experiment.

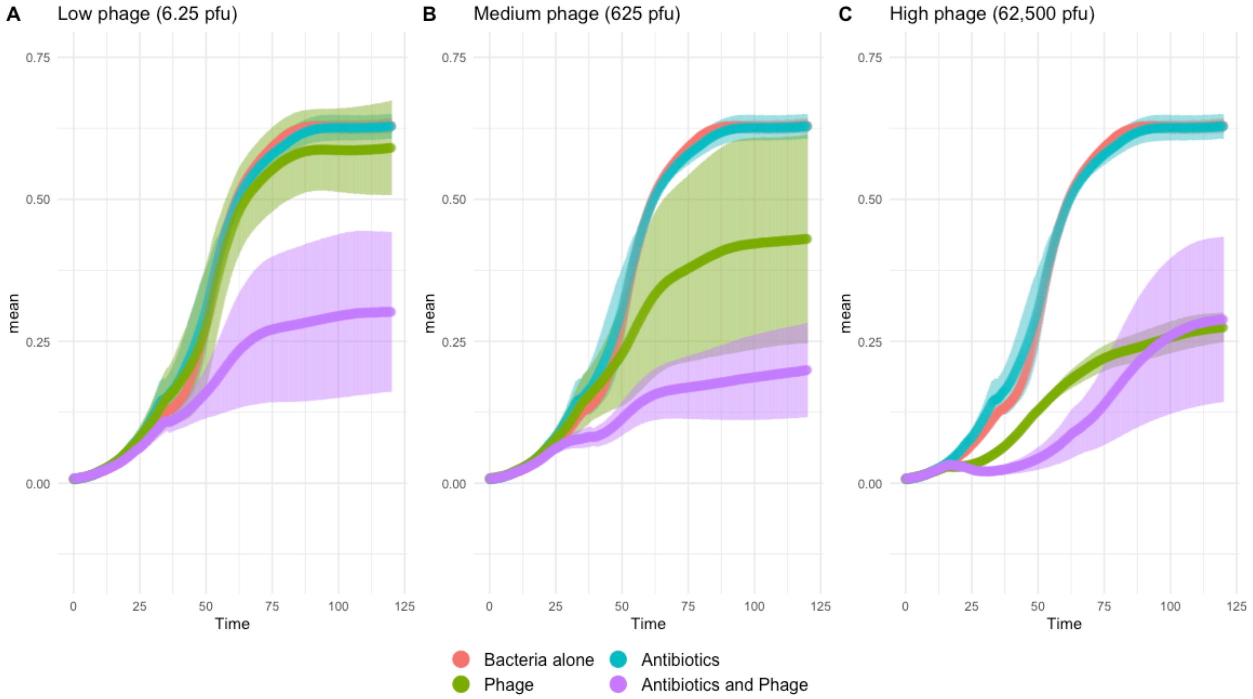


Figure S10: Bacterial growth curves displaying how *Bifidobacterium* is impacted by low dosages of tetracycline (0.1 mg/L) and varying titers of phage.

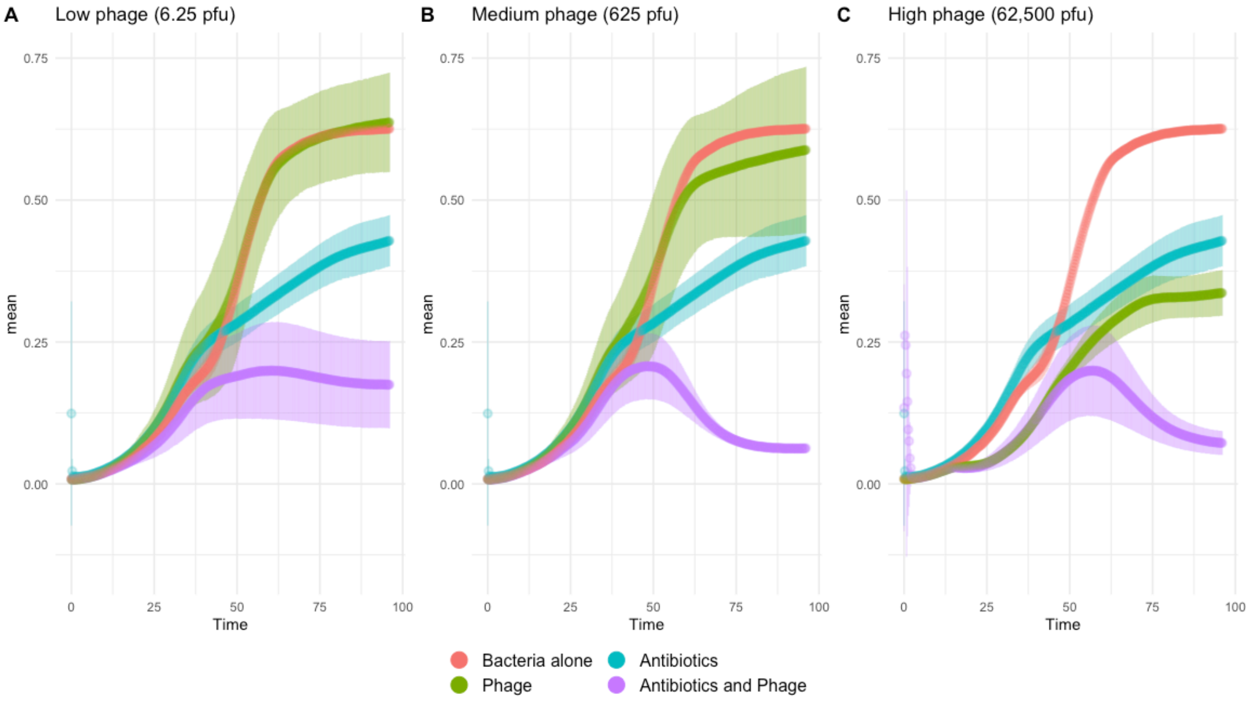


Figure S11: Bacterial growth curves displaying how *Bifidobacterium* is impacted by low dosages of ampicillin (0.1 mg/L) and varying titers of phage.

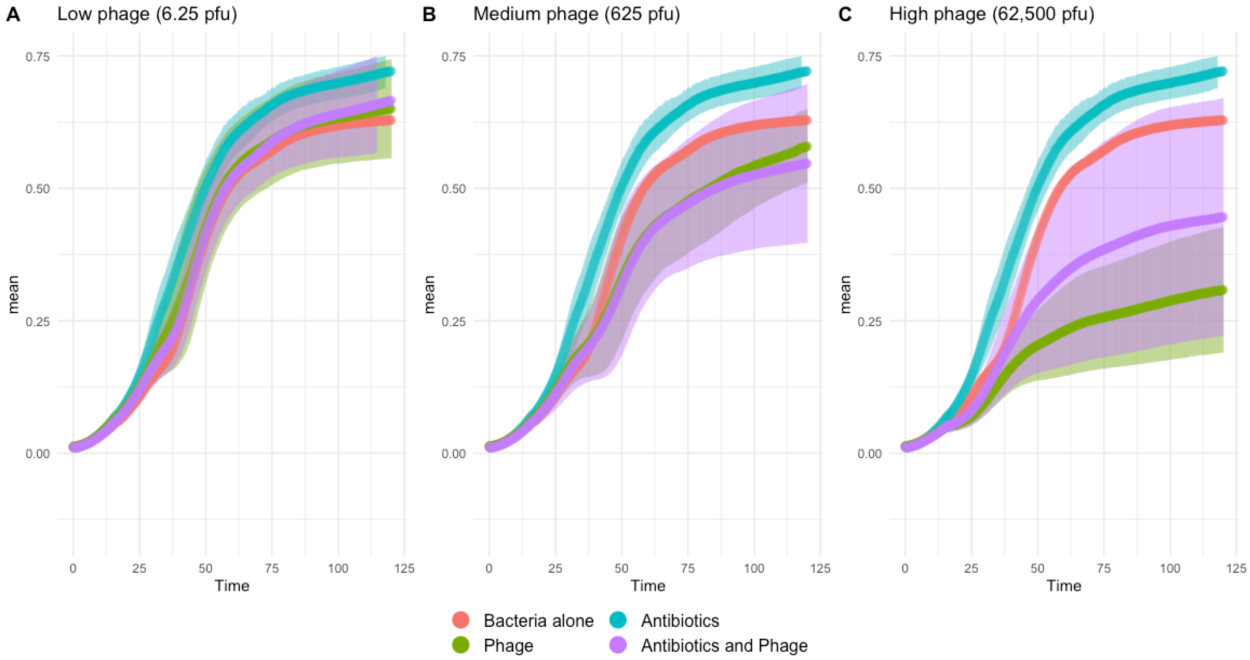


Figure S12: Bacterial growth curves displaying how *Bifidobacterium* is impacted by low dosages of ciprofloxacin (0.1 mg/L) and varying titers of phage.
